# Microdynamics of lamin A Ig-fold domain regulates higher order assembly of the protein

**DOI:** 10.1101/2022.01.27.478013

**Authors:** Chandrayee Mukherjee, Duhita Sengupta, Lakshmi Maganti, M. Mahendar, Dhananjay Bhattacharyya, Kaushik Sengupta

**Affiliations:** Biophysics & Structural Genomics Division, Saha Institute of Nuclear Physics, 1/AF Bidhannagar, Kolkata-700064, West Bengal India, Homi Bhabha National Institute, Mumbai, 400076, India; Computational Science Division, Saha Institute of Nuclear Physics, 1/AF Bidhannagar, Kolkata-700064, West Bengal, India, Homi Bhabha National Institute, Mumbai, 400076, India

## Abstract

Lamins maintain the shape and rigidity of the nucleus in the form of a proteinaceous scaffold underneath the inner nuclear membrane (INM) and provide anchorage to chromatin and other nuclear proteins. Mutations in the human LMNA gene encoding lamin A/C cause about 16 different diseases with distinct phenotypes collectively termed as laminopathies which affect primarily the muscle tissues as well as adipose tissues, neuromuscular junctions and multiple other organs in progeroid syndromes. Lamins contain several domains of which Ig-fold is one of the well characterized and structured domains that harbours many mutations leading to deleterious interactions with other nuclear proteins. In this work, we have elucidated the effects of 3 such mutations namely R453W, W498C and W498R on the dynamics and flexibility of the Ig-fold domain and the consequent effect on the assembly into lamina by live cell imaging, fluorescence correlation spectroscopy (FCS) and molecular dynamics (MD) simulations. From our experimental observations, we concluded that R453W has somewhat increased flexibility compared to the other mutants but lower than that of the wild type. This might be attributed due to an alteration of the salt bridges in the Ig fold domain of the mutant. This is the first report where such an alteration in the full length has been documented by gross changes in diffusional properties as a sequel to a mutation in the Ig fold domain

## Introduction

A and B-type lamins are intermediate filament proteins and form a fibrous network in the nucleoplasmic space beneath the INM of all terminally differentiated metazoan cells except plants (1-4). The lamins constitute the scaffold for the nuclear envelope and facilitate tethering of chromatin and other nuclear proteins (4). Structurally, lamins possess a characteristic tripartite structure unique to intermediate protein family. The central coiled-coil forming α-helical portion of lamins is flanked by a short unstructured ‘head’ domain at its N-terminal and by a longer C-terminal domain (5-7). The C-terminal domain is grossly unstructured except for a tiny island of ∼113 residue long globular motif consisting of nine β strands thus conferring it a classical Immunoglobulin fold (Ig-fold) architecture (8,9). The sandwich-like structure of two β sheets consisting of 5 and 4 β strands respectively are connected by short loops (10). The C-terminal domain harbours binding sites for chromatin (11) as well as other nuclear proteins like PCNA, LAP2α, emerin etc. (12,13). Furthermore, the Ig-fold domain maintains a unique status due to its interaction with PCNA (14), Nup proteins and harbouring a site of interaction with SUMOylated substrates (15).

An enormous number of lamin A/C mutations have been uncovered since the discovery of lamin A/C mutation and its connection with dilated cardiomyopathy (16). Nearly 500 such mutations in lamin A/C have been reported to date (http://www.umd.be/LMNA/) and these mutations cause muscular dystrophies, lipodystrophies, dilated cardiomyopathy and premature aging syndrome affecting multiple organs (10,17,18). On the other hand, lamin B1 duplication was shown to produce autosomal dominant leukodystrophy (19). Interestingly, the recent work from Cristofoli et. al. has linked for the first time the role of missense mutations in LMNB1 with severe autosomal dominant microcephaly (20). This has led to a major impetus of research activities focussing primarily on the mutations of lamin A/C and their consequent effects in cells and tissues. It is important to note that almost 15% of the total number of mutations in lamin A/C map in the Ig-fold domain (10) which implies that almost 75% of the amino acid residues in this domain is mutated to missense mutations. This makes it an interesting domain to delve into and address the structural perturbations associated thereof. Most of these mutations cause Emery Dreifuss Muscular Dystrophy (EDMD), Limb-Girdle Muscular Dystrophy (LDMD) and Dilated Cardiomyopathy (DCM) (http://www.umd.be/LMNA/). We have focussed on three such deleterious mutations R453W, W498C and W498R to check their structural plasticity and the overall effect on the morphology of the nucleus. Out of these 3 mutants, 66 patients have been reported to be afflicted by R453W with an early onset with severe phenotypes. R453W abrogates the two important salt bridges formed by Glu 443 and Glu 444 thereby favouring the transition to unfold rather easily (4). R453W causes AD-EDMD which is characterized by rigidity of the spine, scapular weakness, contractures of elbows and Achilles’ tendons progressive muscle wasting with frequent cardiac involvement (21). W498C was shown to produce LGMD1B in the patients who complained of limb-girdle weakness with special emphasis on pelvic girdle. The tryptophan at 498 is located at the C-terminal end of strand 6 and occupies the pocket formed by the hydrophobic side chains. Y481 on β-strand 5 is another such hotspot that has been also reported to cause LGMD1B phenotype in patients (22). W498C also showed abnormalities in Z-disk arrangement in skeletal muscles with concomitant disorganization of desmin, which is known to connect cytoskeleton to the nuclear surface thus perturbing mechanotransduction from cytoskeleton to nucleoskeleton (23). In retrospect, our group had shown similar perturbation of lamin A-emerin interaction for the mutant R453W and attributed that to modified mechanotransduction in EDMD nuclei (4). Lastly, patients inflicted with W498R were diagnosed with DCM associated with atrioventricular block (AVB) along with AD-EDMD (24).

It must be emphasized at this point that very few studies have been invested to probe residue wise structural fluctuations in the Ig-fold and the consequences thereof. Dialynas et. al. carried out HSQC-TROSY NMR experiments with G449V, N456I, L489P and W514R (25) which was further extended by a detailed investigation of longitudinal/transverse relaxations in W514R complemented by MD simulations (10). In the current work, we have compared local dynamics and diffusional flexibility between R453W, W498C and W498R by FCS and MD simulations. Interestingly enough, we found that R453W exhibited maximum flexural dynamicity over other mutants and the wild type protein which was also evident from its translational motion in the nucleoplasm. We hereby present a novel and interesting insight into the activity of the Ig-fold domain which in turn modulates the diffusional parameters of the full length protein in the nucleoplasm. This has tremendous implication in the rate of formation of the lamin A network and the lamina underneath the INM. Also, this has shed light for the first time on the role of mutant lamin A in the formation of aberrant nuclei thus explaining pathophysiology of the diseases to a greater detail.

## Methods

### Plasmids and constructs

Full length wild type lamin A (henceforth referred to as wt LA), R453W, W498C and W498R were cloned into pEGFP (Clontech) and/or pRFP (Clontech) by techniques as described earlier (6,10,26).

### Cell lines and Transfection

C2C12 (ATCC) was maintained in DMEM (Gibco, USA) supplemented with glucose to a final concentration of 10mM, Penicillin-Streptomycin to a final concentration of 1% and Fetal Bovine Serum (FBS) (HiMedia, India) to a final concentration of 10%. Cells were cultured using standard protocol and were maintained at 37°C under 5% CO_2_. Cells in their early passages were allowed to grow to a maximum of 50-60% confluency in coverslips and 35mm glass bottom dishes. Cells were co-transfected with GFP tagged wt LA / GFP tagged mutated lamin A along with RFP tagged wt LA. 3ug of total DNA was transfected with Lipofectamine 2000 (Invitrogen, USA) (DNA: Lipo2k = 1:1.5) in serum and antibiotic free medium as described earlier (27). Cells were used for fluorescence imaging after 24 hours of transfection.

### Confocal microscopy: Z stack and Live cell imaging

C2C12 were co-transfected with EGFP tagged wt LA/EGFP tagged mutant LA along with RFP tagged wt LA. Transfected cells were fixed and stained as described earlier (27) (26). Slides were visualised under Nikon Eclipse Ti-E microscope with Plan Apo VC 100X oil DIC N2 objective/1.40NA/1.515RI with 4X digital magnification. The images were captured in Nikon Eclipse A1R+ using resonant scanner. During Z stack imaging, step size of 0.15μm was maintained over a z depth of 5μm. Multi line argon ion laser (488 nm laser line for GFP excitation), solid state laser (405nm for DAPI excitation) and a He-Ne laser (for RFP excitation with 561 nm laser line) were used. During live cell imaging, movement of the speckles were visualised by maintaining live cell conditions in CO_2_ independent medium (Gibco, Thermo Fisher, India) for 1 hour with frames captured at intervals of 30 seconds using the Perfect Focus System (PFS) of Nikon Eclipse A1R+ Ti-E confocal system. Z stack images were processed in Ni Element Analysis AR (Version 4.13.00). Pearson Correlation Coefficients (PCC) and van Steensel’s cross-correlation coefficients (CCF) (28) were measured from manually selected regions of interest with the help of JACoP plugin in ImageJ (1.47v). The number of aggregates in the nucleus were also calculated in ImageJ by setting the threshold value to black and white and converting the image to 8 bit and subsequently outlining and calculating the number of particles by the outline tool and the analyze particles tool respectively.

### FCS measurements

The FCS measurements were performed on a Nikon Eclipse TiE Microscope (Nikon, Japan). All the experiments were carried out at 25°C on live C2C12 cells plated on 35mm glass bottomed dish (Nunc, USA) transfected with EGFP tagged wt LA/mutants. Cells were washed off the phenol red of the media and kept in indicator free warm CO_2_ independent media (Gibco, USA) before imaging. The excitation of EGFP was carried out by a solid state diode laser (λ=488 nm) at 50 mW. 60X water immersion objective with N. A=1.2, a pinhole of 1 A.U, and PMA hybrid 40 detectors (Picoquant GmbH, Germany) were used for acquisition. Filter and dichroic mirror were set to 488 nm. An average of 20 AUX ROI for each nucleus was determined. 75-100 nuclei for each variant were captured. All the data were stored in a built-in module of PicoQuant named SymPhoTime 64 (version 1.5). Subsequently the data was plotted and presented by using Microsoft Excel (version 2108).

Due to the free diffusion of molecules in and out of the confocal detection volume, the induced intensity fluctuations by *I*(*t*) each molecule can be presented collectively in the form of Fluorescence autocorrelation function (FAF) or *G*(*τ*) with the help of the following equation,

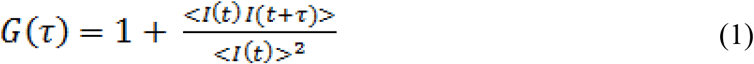

Where, *τ* is the delay time in between two single photon counting method events and < > denote ensemble averages. As the observation volume has a 3D Gaussian profile, the autocorrelation curve fitting for *G*(*τ*) of lamin A proteins were achieved with a 2-component diffusion model (i,1 and 2) in the SymPhoTime 64 (PicoQuant) software, which is given below,

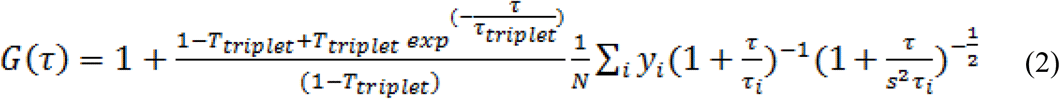

where *N* is the average number of fluorescent molecules in the measurement volume, *T*_*triplet*_ is the average fraction of triplet state of molecules, *τ*_*triplet*_ denotes triplet decay time, *y*_*i*_ and *τ*_*i*_ are the fraction and diffusion time of the component i respectively, *s* is the ratio of *ω*_0_ and z_0_ which are the beam waist and axial radius respectively at the approximated cylindrical measurement volume over which the intensity decay by 1/e^2^ in the lateral and axial directions. We measured the value of *s* in our present confocal setup experimentally from the measurement of the standard dye atto 488 with a well-established diffusion coefficient. The FAFs for Atto-488 (Invitrogen, USA) were fitted using a one component model (i,1) and the determined value of *s* (*s=* z_0_ /*ω* _0_) was fixed for fitting the FAFs obtained from the live cell measurements (*ω*_0_=0.33 μm and z_0_=1.65 μm). Using the following equation,

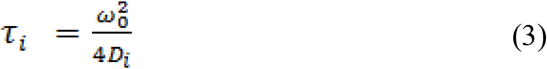

the time of diffusion of component i could be calculated since we determined a fixed value for *ω*_0_ and the translational diffusion coefficient or the *D*_*i*_ for the experiments could be determined from the FAF curve fittings from the live cell measurements.

The diffusion of a non-interacting presumably spherical protein can be defined by the Stokes-Einstein equation,

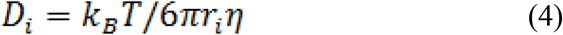

In the above equation, *k*_*B*_ is the Boltzmann constant, T is the absolute temperature, *r*_*i*_ is the hydrodynamic radius of the spheroidal molecule and *η* is the solvent fluid-phase viscosity.

### Western Blot

Cells were harvested post 24 hours of transfection for western blot. The pellets were washed, with sterile 1X PBS, and stored in -80°C till further use. For lysis, the pellet was treated with Mammalian Protein Extraction Reagent (M-PER) (Thermo Scientific, USA) along with 1X protease inhibitor cocktail (PIC). The lysates were stored at -80°C. Protein concentrations in cell lysates were measured using Bradford assay (Biorad, USA). The standard curve was prepared with different concentrations of Bovine Serum Albumin protein. Absorbance was measured at 595 nm and experiments were performed in triplicates. Equal amount of protein sample and 2X Laemmli buffers were mixed to reach the final concentration of 1X Laemmli buffer. The mixture was boiled in a 100°C dry bath for 5 minutes prior to gel loading. Proteins were electrophoretically separated using 10% SDS gel and transferred onto nitrocellulose membrane (Millipore, USA). The membranes were blocked with 5% Non-Fat Dry milk in PBST (1X PBS with 0.1%Tween-20) for 1 hour at room temperature. Primary antibodies used in this study were Rabbit Anti Lamin A (L1293 Sigma Aldrich, USA) in 1:500 and Mouse Anti β Actin (Sigma Aldrich, USA) in 1:1000 dilutions, Antibodies were diluted in blocking buffer and membranes were incubated overnight at 4°C with primary antibodies. Secondary antibody dilution was 1:400 for both anti-mouse and anti-rabbit IgG conjugated to Horse Radish Peroxidase (Santacruz, USA). Membranes were incubated with secondary antibody for 2 hours at room temperature. Following incubation and washing, blots were developed using Enhanced Chemiluminescent (ECL) substrate (Thermo Scientific, USA). An X-ray film was exposed to the signal and was developed using developer and fixer solution (Kodak, USA). After the film dried up, bands were scanned and analysed.

### Molecular dynamics simulations

All-atom MD simulations were carried out for 500 ns under explicit water solvent conditions for WT and its mutants R453W, W498C and W498R. MD simulations were performed at 300 K at the molecular mechanics level implemented in the GROMACS 5.1.3 software package (29) using the CHARMM37 force field (30). Initial structure of the native protein was taken from PDB (31) with PDB ID 1IFR (9). All systems were soaked in a cubic box of water molecules with dimensions such that the box ends are at least 10 Å away from the solute edges. Further, the systems were subsequently immersed in a box having TIP3P water molecules (32). Na^+^ and Cl^−^ ions were added further in the systems for neutralizing. All the systems were minimized using 1,500 steps of steepest descent algorithm to ensure that the system has no steric clashes, inappropriate geometry or structural distortions. All systems were equilibrated at a constant temperature of 300 K by utilizing the two-step ensemble process (NVT and NPT) for 100 ps. Initially, the Berendsen thermostat (33) with no pressure coupling was employed for the NVT (i.e., constant number of particles, volume, and temperature) canonical ensemble, and then we use the Parrinello–Rahman (34) method with pressure of 1 bar (P) for the NPT ensemble (i.e., constant particle number, pressure, and temperature). The LINCS algorithm (35) was employed to constrain bond lengths. Long-range electrostatic interactions were treated with particle mesh Ewald summation method (36) and a cut off of 12 Å was used for short-range interactions. The final simulations were performed for each system for 500 ns where leap-frog integrator was applied for the time evolution of trajectories. The integration time step was set to 2 fs and the trajectories were analysed using built-in programs of GROMACS software package (29). The electrostatic surface potential of the proteins was calculated using the APBS plugin tool of PyMOL.

### Analysis of MD Trajectories

All the trajectory files were analyzed using trajectory analysis module embedded in the GROMACS simulation package, PyMOL (The PyMOL Molecular Graphics System, Version 2.0 Schrödinger, LLC.) and Visual Molecular Dynamics (VMD) software (37). To assess the convergence of the MD simulations, root mean square deviations (RMSDs) were calculated. The energy minimised structures for MD were used as the reference structures in RMSD calculations. In addition, root mean square fluctuations (RMSFs) were calculated to evaluate the structural flexibilities of proteins. The RMSFs were calculated using the final 100-500 ns trajectories of equilibration MD simulations. Principal component analysis (PCA) is based on diagonalization of the covariance matrix of atomic fluctuations to obtain orthogonal eigenvectors (also called principal modes) and the corresponding eigenvalues. PCA was also performed with the GROMACS software package.

### Principal component analysis

Principal components analysis (PCA) is an effective tool for extracting the essential information from MD trajectories by filtering global slow motions from local fast motions (25,38-40). PCA is based on diagonalization of the covariance matrix of atomic fluctuations to obtain orthogonal eigenvectors (also called principal modes) and the corresponding eigenvalues. The eigenvalue indicates the magnitude in the direction of the principal mode. The principal components (PCs) are the projections of a trajectory on the principal modes, of which usually the first few ones are largely responsible for the most important motions. The PCA method has been widely and successfully used to investigate the motion tendencies of proteins. In this work, to detect the conformational change of wild type (WT), R453W, W498C and W498R mutant, PCA was also performed with the GROMACS software package (29).

### Dynamic Cross-Correlation Map (DCCM)

DCCM analysis was performed to evaluate the dynamic correlations of domains and compare the correlation matrix across all Cα atoms for all the systems. The correlation coefficient *C*_*ij*_ between two atoms i and j during the course of the simulation trajectory is defined as (41):

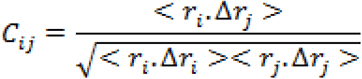

where displacement vectors Δ*r*_*i*_ or Δ*r*_*j*_ are calculated by subtracting the instantaneous position of *i*^th^ or *j*^th^ atoms with its average position. *C*_*ij*_> 0 represents the positively correlated motions between the *i*^th^ and *j*^th^ atom and *C*_*ij*_< 0 represents the negatively correlated motions between them. Dynamic cross-correlation map (DCCM) analysis was performed using Bio3D (42).

## Results

### Abnormal assembly and movement patterns of lamin A mutants

We had shown earlier that overexpression of mutant lamin A in C2C12 cells led to aggregation throughout the nucleoplasm (6,10,19,43,44), irrespective of the position of the mutation along the length of the protein. In our present work, we have analysed the differential pattern of assembly of wild type lamin A protein and its mutants (R453W, W498C, W498R) inside the cell. The point mutations under study are localized in the Ig-fold domain and have been depicted in a cartoon (Figure 1A). Considering the heterozygosity of the mutants, we have co-transfected the cell lines with EGFP tagged lamin A constructs (wt/mutant) and RFP tagged wt LA as shown earlier (45,46) and performed Z stack imaging of the fixed cell samples to analyse the three-dimensional distribution of the fluorescent proteins throughout the entire volume of the nucleus. The rationale behind this approach is to mimic the effect of the mutant allele on the wild type and its consequence on the formation of lamina. In the first case where the cells were transfected with both EGFP and RFP tagged wt LA proteins, we observed uniform and intact lamina formation from the 3D distribution of LA which recapitulated the usual characteristic organization of endogenous wt LA underneath the INM. The distribution of LA was continuous and uniform across the entire Z depth (Figure 1B). However, when the cells were co-transfected with each of EGFP tagged mutant LA (R453W/W498R/W498C) and RFP tagged wt LA, overexpression mediated aggregation of mutant proteins could be observed concomitantly with the formation of a broken nuclear rim (Figure 1B). The discontinuity in the nuclear rim and abnormal punctate distribution of LA across the Z depth was noticed due to aggregate formation in the mutants (Figure 1B). Comparatively stable and higher number of aggregates were found in lamin R453W (Figure 1B). 90-95% of the positively transfected cells showed a similar phenotype for each of the individual mutants. The RFP tagged wt LA was always found to be sequestered by the mutant aggregates (Figure 2A). Ectopic expression of the wt LA along with the mutant protein confirmed the sequestration event which was evident from co-localization of the two. Higher numbers of such aggregates were observed for lamin R453W with a PCC value of 0.98 followed by W498C and W498R with PCC values 0.95 and 0.84 respectively whereas the co-expression of the wt LA with RFP and GFP fluorophore resulted in a few tiny puncta with a PCC value of 0.69 which is common in all ectopic expression scenario (Figure 2A). Higher PCC values in the mutant proteins indicate a greater correlative variation of GFP tagged mutant lamin A along with RFP tagged wt LA. The fact that the phenotypes observed for the mutants didn’t result from expression artefact but were true effects due to the mutations, was corroborated from the expression profile of the proteins in western blot (Figure 2B). On quantification of the number of visible aggregates in the nucleoplasm, largest number of aggregates were observed for R453W (∼25) while lowest for wt LA (not more than 5) (Figure 2C). To summarize these observations, we conclude that these mutations showed a preponderance of non-specific aggregation over stepwise assembly into an intact lamina and the aggregates of varying sizes acted as “sequestration sinks” which are in stark contrast to the wild type which predominantly formed an intact lamina along with some tiny aggregates. In R453W aggregates, greater spatial correlation of red and green fluorescent probes was observed with respect to the other two mutants depicting higher proportional co-distribution of the two fluorescent proteins. The van Steensel’s cross correlation function suggests how the PCC alters as a function of the shift of red image pixel to green. Peaks at the centre in CCF plots for all three mutants as well as WT variety depict that the fluorescent signals from the two channels in the mutants and the wild type were all positively correlated but to different extents. We also observed differential mobility of the mutant aggregates with infinitesimally small movements for the wt LA (Supplementary video S1). Since there were minute or no translational artefacts of living cells on the glass plate over 3600 seconds, the observations were not processed further to draw any significant conclusions (Supplementary video S1). To summarize the results obtained, we invoke a model to explain the abnormal ability of the mutants to form non-specific aggregates following a course of random collisions rather than exhibiting a slow, systematic and stepwise assembly to a functional lamina of giant polymeric structure and almost zero dynamicity. This prompted us to analyse the diffusional flexibility of the full-length proteins in vivo by fluorescence correlation spectroscopy and later on to study the mobility of the very specific domain harbouring the mutations in a more precise manner by molecular dynamic simulations.

**Figure 1.**
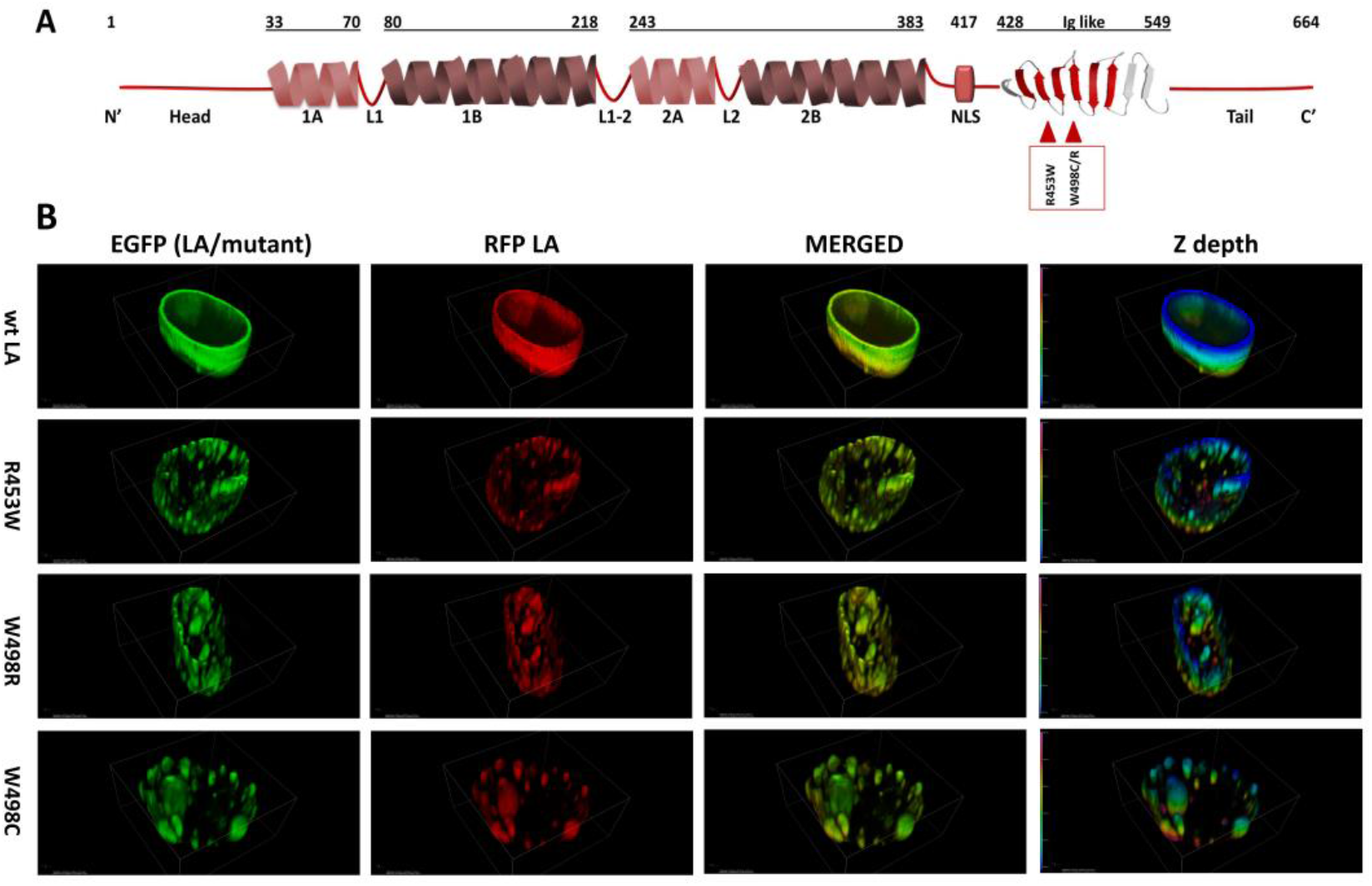
Abnormal nuclear morphology induced by lamin A mutants. **(A**) Schematic of lamin A protein indicating the location of point mutations on the Ig-fold studied in this repor**t**. (**B)** C2C12 cells were transiently co-transfected with RFP wt LA and GFP WT/ R453WLA/W498CLA/W498RLA. Post 24 hours of transfection, cells were fixed and stained and each nucleus was scanned for a Z depth of 5μm 3D reconstitution of images along Z axis showed a continuous and uniform distribution of lamin A in wt LA whereas all three mutants produced punctate lamina with irregular distribution across the z depth. Comparatively stable and higher number of aggregates were found in R453W. The last panel exhibits the psedu-colored 3D z depth rendered basal, apical and side views of the nuclear rim indicating an uneven distribution of the aggregates and differences in their size and pattern.

**Figure 2.**
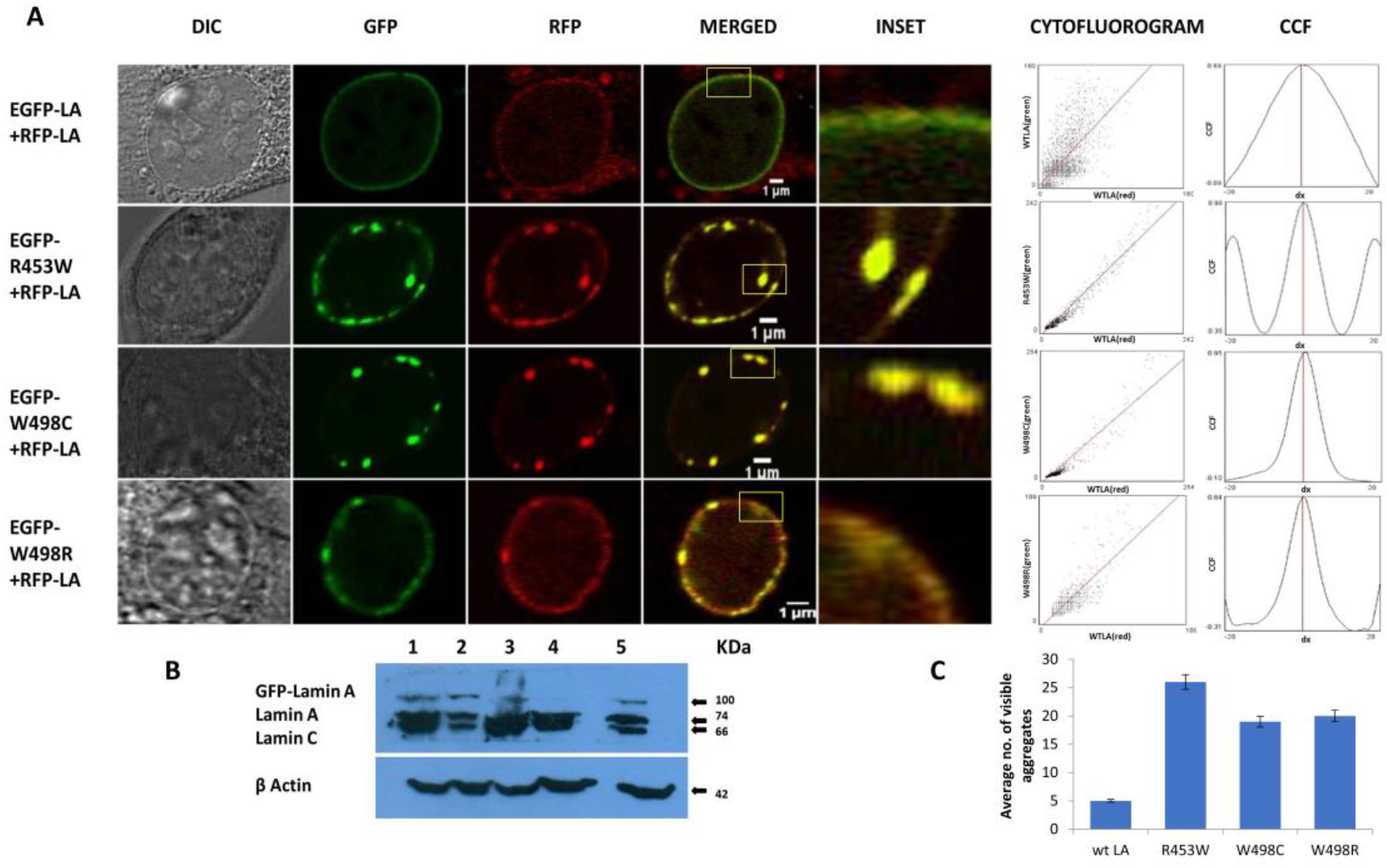
Sequestration of RFP LA by the mutant aggregates: **(A**)Live cells were imaged for 1 hour. One frame from wt LA and each of the mutants has been selected to show the pattern of speckle formation. Insets of the panels are showing sequestration of GFP tagged WT and mutant lamin A proteins with RFP tagged wt LA in forms of yellow aggregates. PCC values from respective cytofluorograms and van Steensel’s cross co relation functions are showing positive correlations in all four co transfection scenario and higher PCC values in the mutant counterparts are indication of stronger association of red and green probes in the yellow aggregates delineated with the ROI shown in the inset. Magnification: 100x. Scale Bar: 1 um. (**B)** Western blot showing prominent band of lamin A tagged with EGFP (100kD) along with bands for endogenous lamin A/C (74kD/66kD). Lysates from transfected cells were immunoblotted using Anti Lamin A antibody to check for EGFP LA expressions. **1-**R453W, **2-**W498C, **3-**W498R, **4-**Non-Transfected, **5-**wt LA. ß Actin (42kD) is used as loading control. (**C**) The average number of visible aggregates observed in the nucleoplasm for wt LA and the mutants. Error bars indicate percentage error.

### Diffusional landscape of mutant lamin A proteins in the nucleoplasms

It is well established from earlier studies that the nuclear lamina has lamin proteins having little or no mobilities but the soluble fraction of lamin A proteins are mobile and the diffusion parameters have also been well characterized earlier by FRAP (47) and subsequently with the help of FCS in a number of studies(45,48,49). Therefore, taking cues from those studies, we have compared the diffusional motions of full length lamin A and its 3 mutants by employing FCS based microscopy technique whereby we get information on their diffusivity, dynamics and direct measure of stoichiometry with almost single molecule sensitivity (Figure 3) in an infinitesimally small volume of femtolitres. Fluorescence intensity fluctuations in the nucleoplasm of transiently transfected live C2C12 cells expressing GFP tagged wild type and mutant Lamin A were recorded individually at 24-48 h after transfection. A typical C2C12 cell expressing wild type lamin A is shown (Figure 3A). Cells expressing GFP-tagged proteins were localized by confocal imaging and the confocal volume was assigned at arbitrary locations in the nucleoplasm, carefully avoiding the aggregated proteins of much larger brightness in case of the mutants. Measuring time was set to 10s as described earlier (50) which was appropriate to obtain reliable autocorrelation functions. A representative photon count trace, measured with FCS, from EGFP-LA in nucleoplasm (Figure 3B) is shown. For comparison of the diffusion parameters, the amplitude of G(τ) (G (0)-1) was normalized to 1. The diffusion constants of the full length lamin wt LA and the mutant LAs are summarized in the table (Figure 3C). Faster diffusion times corresponding to the FAFs of wt LA, R453W, W498C and W498R turned out to be 8.4±0.54, 12.8±0.11, 5.8±0.79 and 7.6±0.63 milliseconds respectively (Figure 3D). On the other hand, the slower diffusion times didn’t vary too much. It must be emphasized that the two diffusion times obtained from the two-component analysis correspond to a slow mobile phase and a fast mobile phase as described earlier(45,48). W498C exhibited slightly faster diffusivity followed by the wild type and W498R which were almost similar whereas the slowest value was recorded for R453W (Figure 3E, F and G). This points to the fact that R453W diffuses rather slowly which could be due to an increase in its bulk which might result from elevated aggregation propensity. We conjectured that this differential diffusional dynamics of the full length protein could be a direct consequence of the microdynamics of the Ig fold domain which harbours the mutations. To test this hypothesis, we performed MD simulations of the Ig fold domain subsequently.

**Figure 3:**
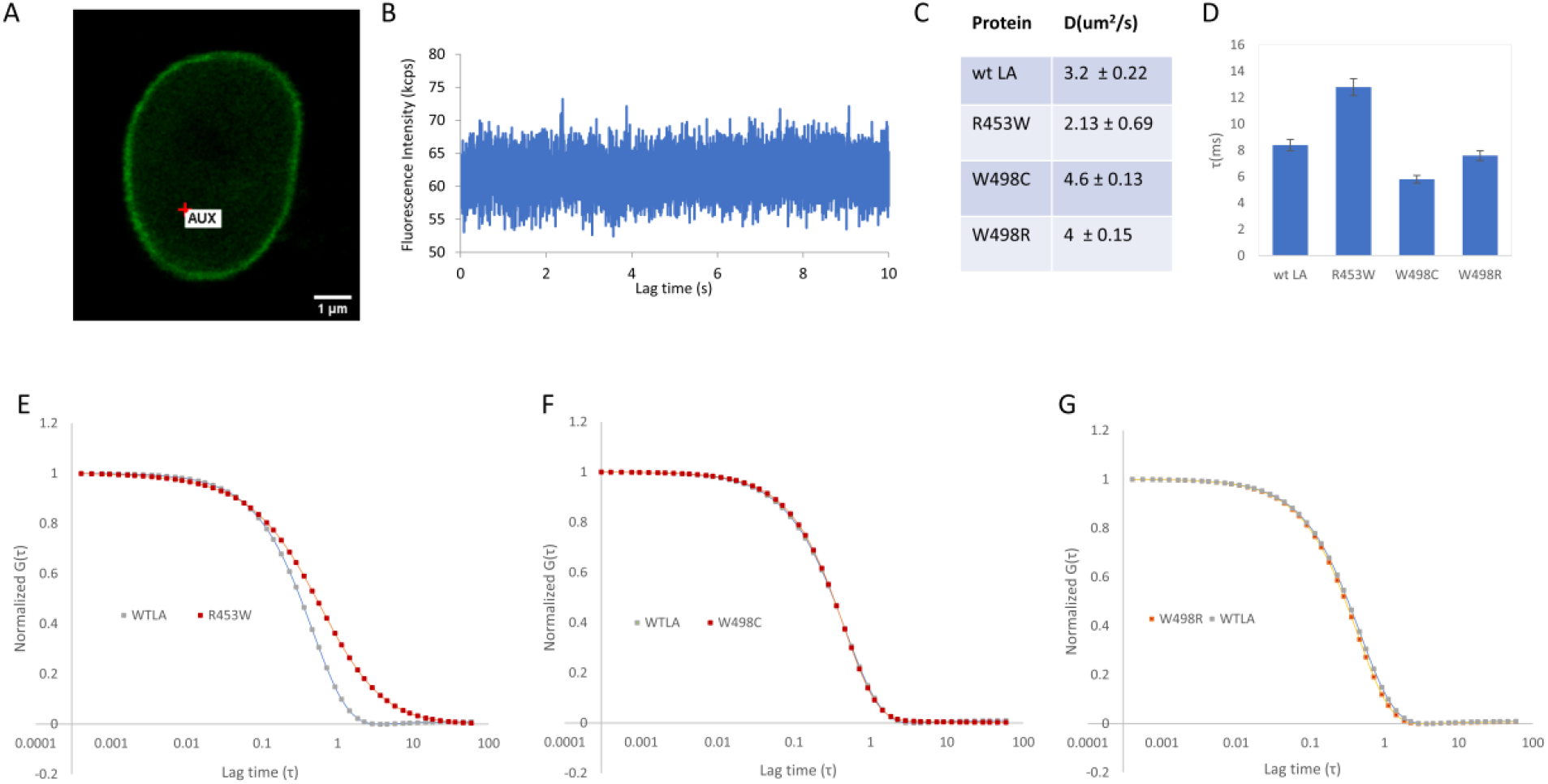
Study of the nucleopalsmic diffusional mobility of wild type and mutant full length proteins in vivo by Fluorescence correlation spectroscopy: The differential mopbilities of wild type GFP lamin A (wt LA), R453W, W498R and W498C in C2C12 cells were analysed by FCS. (A) A representative image of a nucleus expressing wild type GFP lamin A during FCS measurements. The red point indicates the ROI where the measurements were obtained. Scale bar: 1μm. (B) Representative fluorescence fluctuation of GFP-LA in the nucleoplasm has been plotted as a function of time. (C) Table displays the diffusion coefficients of mutant and wild type full length lamin A proteins in live cells as determined by FCS. (D) The average diffusion times corresponding to the diffusion coefficients of the proteins. Error bar indicates percentage error (E, F and G). Averaged fluorescence autocorrelation curves were calculated from the fluorescence intensity fluctuations and plotted as a function of time in seconds. G(τ) is the fluorescence autocorrelation function. For comparing the diffusional mobility, all curves were normalised to the amplitude G(τ)=1 at 0 seconds.

### Association and dynamics of the Ig-fold

We analysed the electrostatic surface potential and performed MD simulations for 500 ns on the wt and the three mutant Ig-fold domains. The root mean square deviation (RMSD) was monitored during the simulation time to investigate the protein stability. The RMSD values converged very well in all the systems, within 100ns, thus the MD trajectories from 100–500 ns were chosen for further conformational analyses (Supplementary Figure 1). Mean values of RMSD of the wt, R453W, W498C and W498R system were approximately 1.81 ∼ 2.34 Å, indicating that the 3D structure of the Ig-folds were stable in all the cases. The structures after 500 ns MD simulations, along with their electrostatic potential surfaces are shown in Figure 4, which indicates that the mutations didn’t result in gross structural perturbations of the protein. The Ig-fold was maintained in each system with its signature β-sheet sandwich. The electrostatic potentials, as calculated by APBS toolkit of PyMOL, however, indicated major changes in polarity of the protein upon most of the mutations. The wt Ig-fold shows prominent negative and positive patches near N-terminal (residue 436) and the long loop near 512 which faded out in the R453W and W498C. In case of W498R, another region near N & C termini assumed a highly positive patch. This might lead to an increased tendency of non-specific association for the mutants resulting into aggregates/speckles. However, the root mean square fluctuations (RMSF) of all the systems indicated that the amplitude of atom displacements was different among the wt and the mutants (Figure 5). Most of the residues adopting β-sheet conformations exhibited low RMSF whereas the loop regions showed larger fluctuations with wide variations. For instance, the loop region between the two parallel stranded β-strands (6^th^ and 7^th^, residues 500 to 515) showed higher fluctuation for most of the mutants and the wild type. The R453W mutant showed highest fluctuations for residues 475 and 525, which remained close to each other in Euclidian space. It may be noted that the site of mutation (453) is quite far from the regions of high fluctuation. Therefore we can speculated that there might be some long-range effect in the otherwise compact protein. The positively charged side-chain of Arg-453 protrudes out towards solvent for the native protein, which is mutated to hydrophobic Trp, by single nucleotide alteration in the genomic sequence, in the diseased form, which may be accommodated by different ways. The protein might destabilize the third β-strand by pushing the Trp side-chain towards core of the β-sandwich or assume multimeric form by association with other proteins. In our computational study, we cannot simulate protein-protein association but we observed the Trp side-chain remain exposed through long range perturbation.

**Figure 4.**
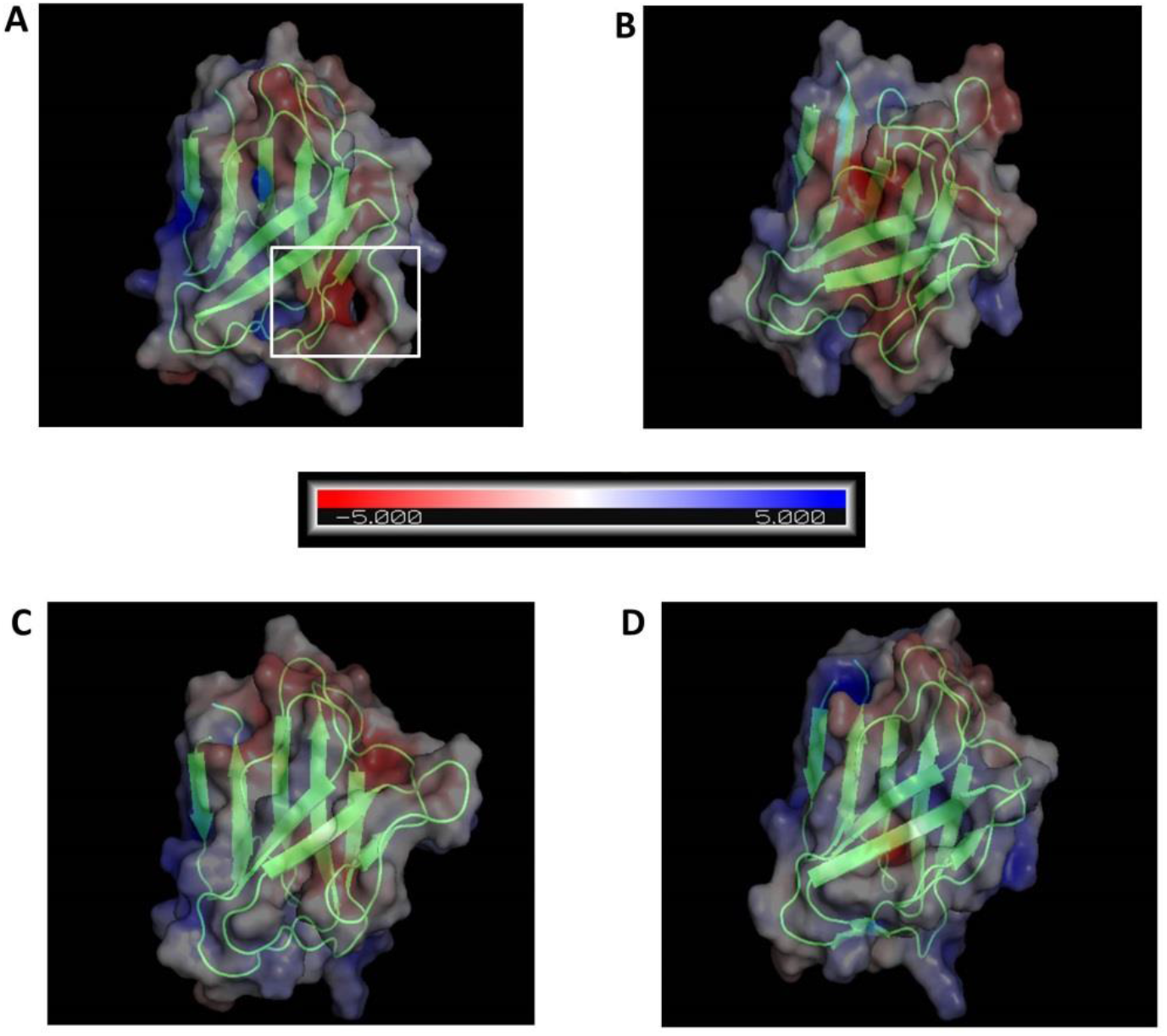
Electrostatic surface potential along with secondary structure of each of the four mutants. **(A)** WT, **(B)** R453W, **(C)** W498C and **(D)** W498R, showing conservation of secondary structure but reduction in polarity for most of the mutants. Scale bar denotes transition from red to blue where red indicates negative potential and blue indicates positive potential.

**Figure 5.**
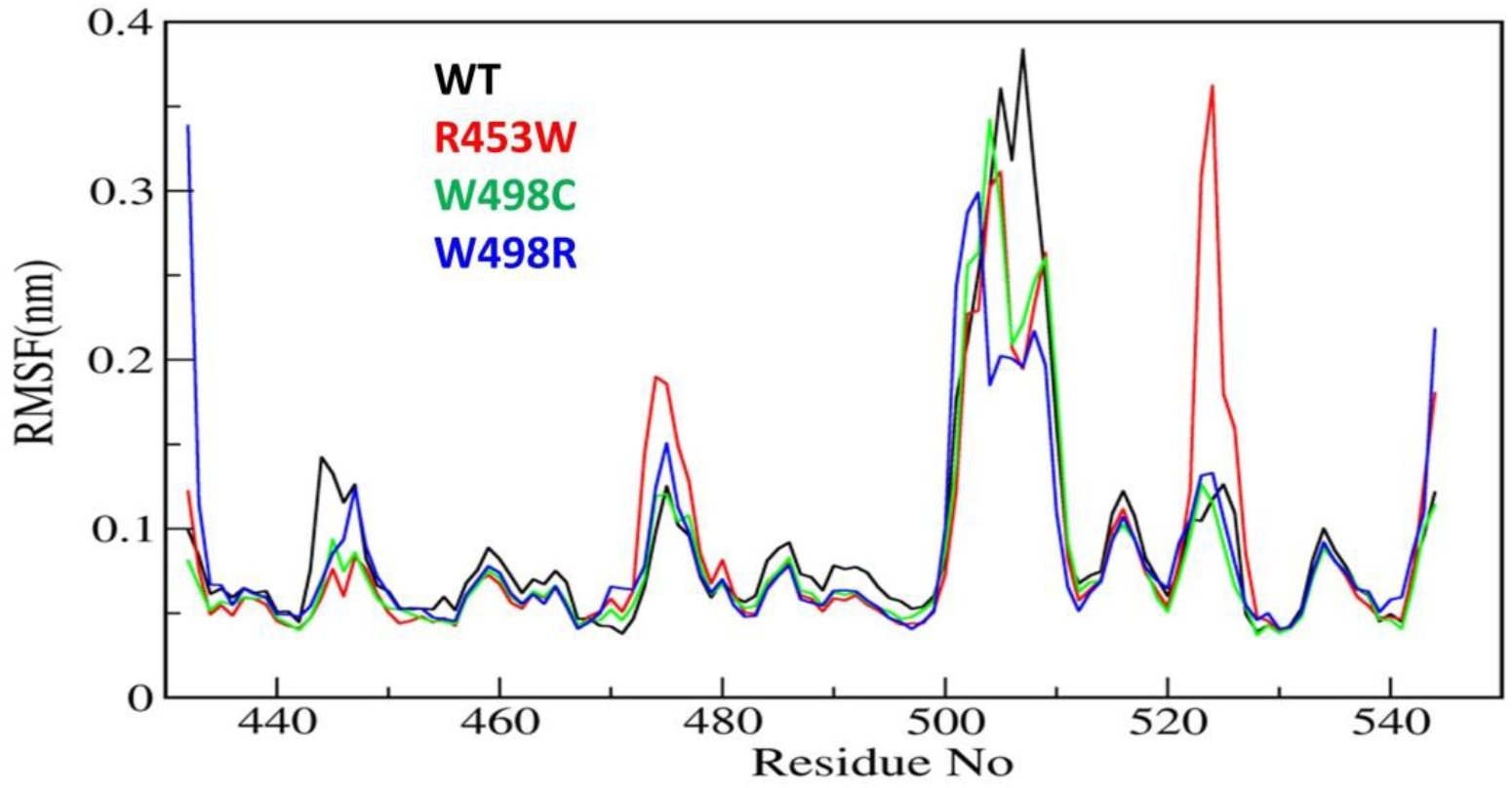
Root Mean Square Fluctuation (RMSF) of Ig-fold proteins. The main difference in structure of the wild type and R453W mutant protein is at the region involving the residues 475 and 525. Color coding-Black: WT, Red: R453W, Green: W498C, Blue: W498R.

### Backbone & secondary structural analysis

It is a well documented fact that movements of important amino acids within particular secondary structural motifs may alter secondary structures of the whole protein to a certain extent. We have analysed alterations of secondary structures of each amino acid in each mutant using DSSP algorithm (51) to understand such alterations of secondary structures (Figure 6). Most of the important differences in the backbone flexibility are mainly localized in the vicinity of the mutated residue. In native, the residues around 512 existed as “bend” conformation till the end of the simulation whereas in all mutants this region retained “turns” conformation only. Between the residues 492-495, the wt showed “turn” with traces of “bend”, whereas in the mutants primarily “turn” conformation was mainly prevalent. In the 455 residue region, β sheet conformation was observed throughout the simulation time in wt, whereas in mutant “turn” was getting disrupted and attained coil conformation. To summarize this part, very slight changes in secondary structure conformation was observed in the WT and the mutants. A few residues near Ser-525 of the R453W mutant showed alteration of structural features. These alterations are results of multimodal distributions of φ and ψ torsion angles. In the R453W mutant, Glu-443 showed unusual fluctuations in its φ and ψ torsion angles due to abrogation of the salt bridge between Glu-443 and Arg-453 present in the wild type.

**Figure 6.**
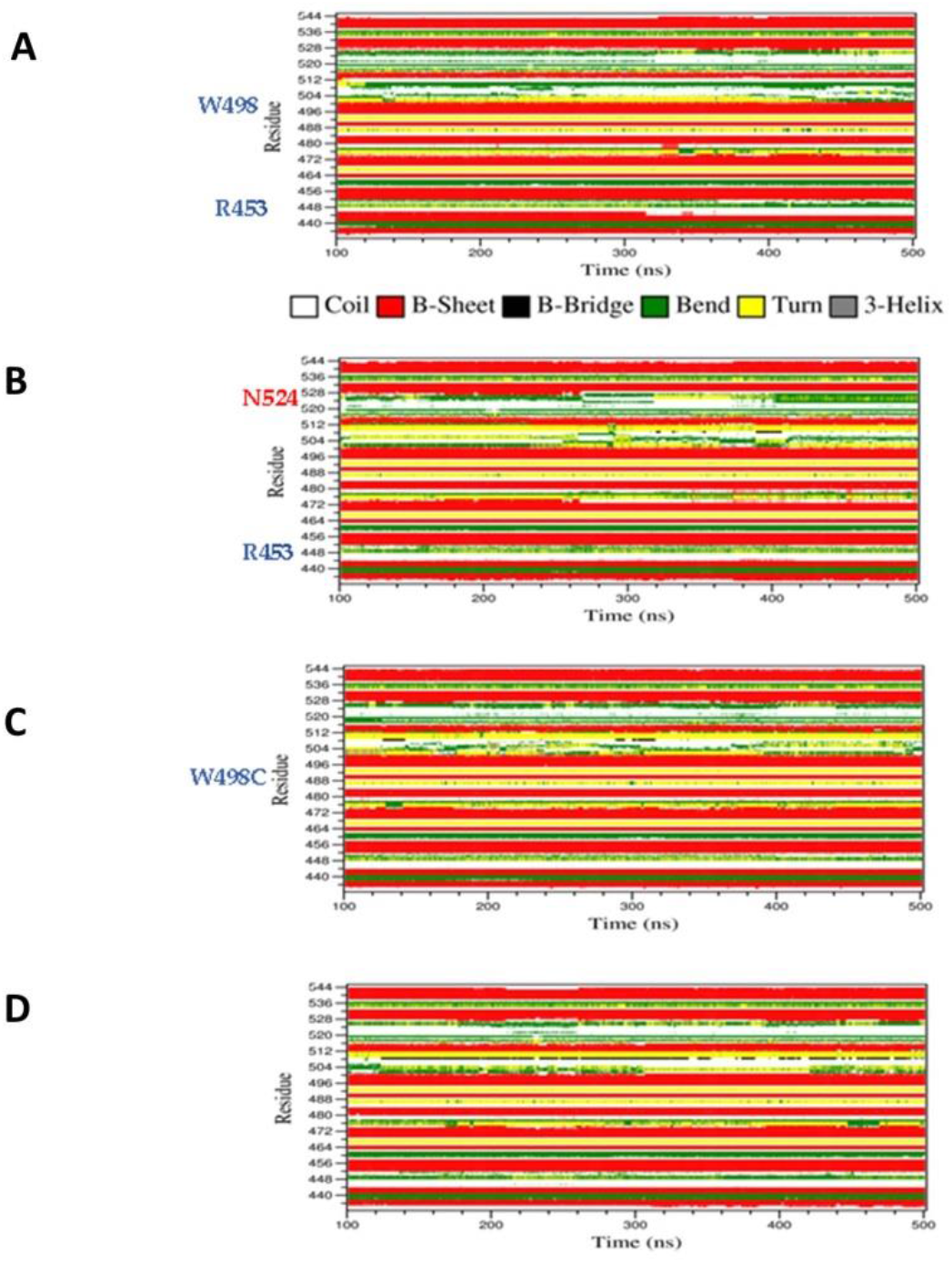
Spatio-temporal variation of secondary structures of the proteins during simulation. **(A)** WT, **(B)** R453W, **(C)** W498C and **(D)** W498R.

### Mapping the dynamics by PCA

Principal components analysis (PCA) is an effective tool for extracting the essential information from MD trajectories by filtering global slow motions from local fast motions (25,38-40). The eigenvalue indicates the magnitude in the direction of the principal mode. The principal components (PCs) are the projections of a trajectory on the principal modes, of which usually the first few ones are largely responsible for the most important motions. The PCA method has been widely and successfully used to investigate the motion tendencies of proteins (25,38-40). In this work, PCA analyses were performed to extract the essential dynamical motions by filtering global slow motions from the fast motions for WT, R453W, W498C and W498R systems. The movements along the first principal component (PC1) from PCA analysis is presented in Supplementary Figure 2. The length of cone represents the magnitude of motion and the pointing of the arrow shows the direction. In all the systems fluctuations were observed near the T505 and P506 regions. In R453W, pronounced fluctuations were observed near the G523 & N524 regions when compared to other systems. However, the motions in other regions were absent. Interestingly, it was found that the three proteins, namely wt, W498C and W498R showed similar features in the movements of amino acids near residue numbers 505 (in the loop between 6^th^ and 7^th^ β-strands) with respect to the N and C-terminal regions of the domain. The R453W, however, showed opposite motions of the same residues. This might be again due to loss of the salt-bridge between R453 and E443 side chains.

### Dynamic Cross-Correlation Matrix (DCCM)

As we found altered motions of some of the residues far from the points of mutations, it appears that the structural cue of the mutations might be transferred through the secondary structural elements by some consorted motions. We have thus characterized the correlated motions between residues of wt LA and the mutant proteins (Figure 7). The dynamic cross-correlation matrix was used to identify the correlated motions and communications between different residues. Correlated motions were extracted in domains of the structures, and normalized to [-1,1]. The negative values of DCCM represent the functional displacement of Cα atoms in the opposite direction and vice-versa. The DCCM map is symmetrical, and the diagonal values, corresponding to correlation between every i^th^ and (i+1)^th^ amino acid, represent the maximum correlation. We hypothesize that the close emergence of positive and negative correlated motions makes the domain less stable. We observed high anti-correlation in the native structure compared to mutant structures indicating a more compact structure of the wild type. The positive correlation between residues of β-strands 3^rd^, 6^th^ and those of 4^th^, 5^th^ indicates a passage of dynamics across the two β-sheet planes. It may be noted that the loop regions consisting of residues 457-467 and 483-493 have some β-sheet property, indicating these pairs of strands had some continuity. This mode of extended β-sheet or β-helix like property was prominently visible in the wild type.

**Figure 7.**
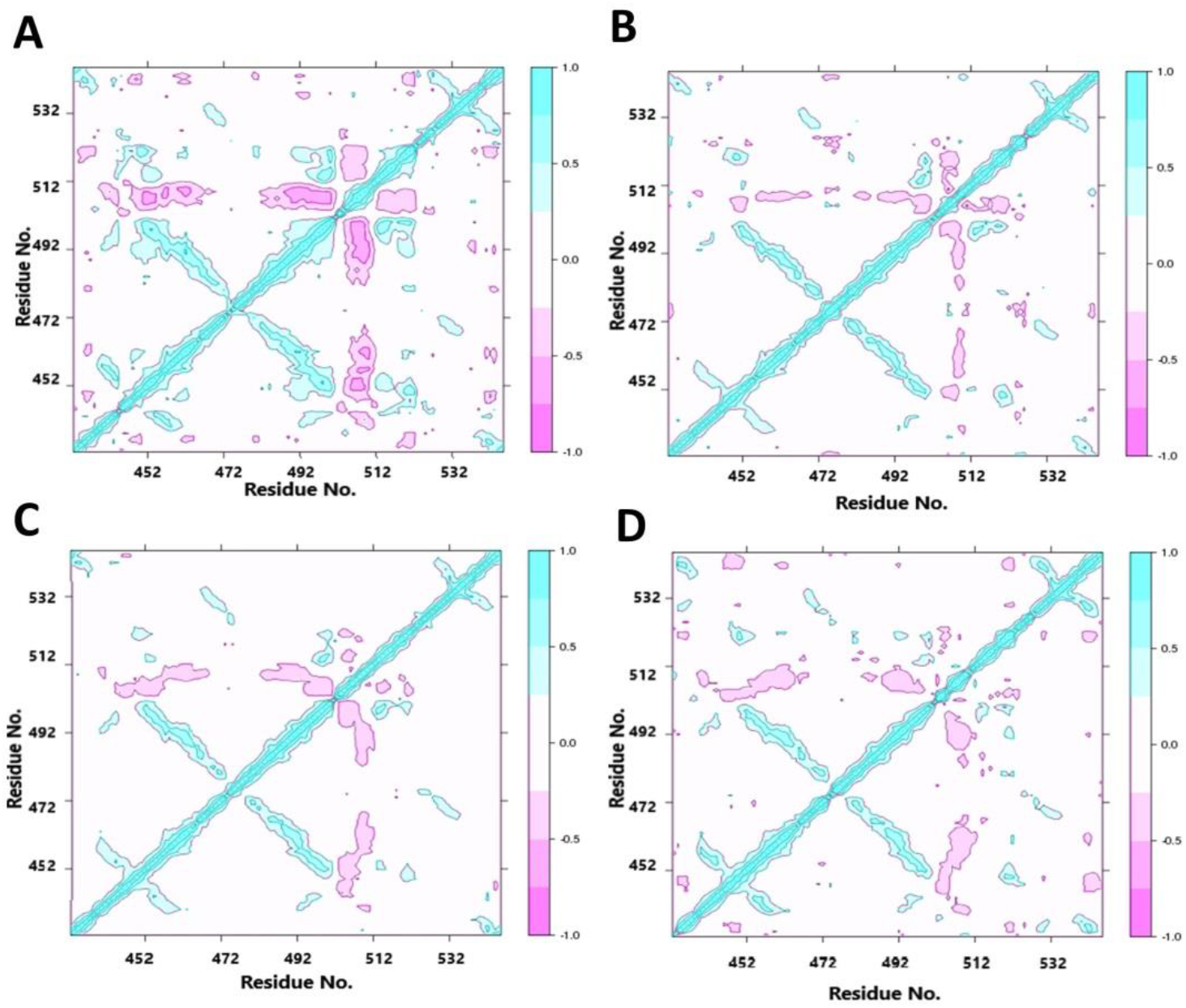
Dynamic cross-correlation matrices of the proteins. (**A**) WT, (**B**) R453W, **(C)** W498C and **(D)** W498R proteins. Cyan represents a positive correlation and pink represents a negative correlation. The color gradient represents the gradual decrease in the correlation

## Discussions

The Ig-fold domain of lamin A has important implications in deciphering the pathophysiology of laminopathies as it is a mutational hotspot. The availability of X-ray and NMR structures (8,9) has spearheaded active research into structural perturbation upon the introduction of mutations. Ig-fold domain of consists of nine β-strands. The sheet of five β strands, 1, 4, 5, 8, and 9 make an angle of 45° with another sheet comprising of 4 strands 2, 3, 6, and 7 thus making up a characteristic β sandwich structure (4,8,9). Previous reports had proven the abolition of beta bulge due to the mutation R453W where the salt bridges with Glu 443 and Glu 444 are abrogated thus facilitating the route for unfolding (4). The same was also proven by steered molecular dynamics and single molecular force spectroscopy. As a consequence of this structural alteration, we had also observed the aberrant self-association behaviour of the mutant protein due to its electronegative surface potential (4). In addition to these earlier findings, we have made in-depth analyses of this protein along with two other mutants W498R & W498C both inside the cell and *in silico* by MD simulations on the Ig-fold domain. We have visualized formation of misshapen and aberrant nuclei in the case of these mutants when we overexpressed the proteins in cells. Formation of normal, continuous lamina was abrogated by aggregates of varying sizes and numbers. Ectopic expression of R453W produced smaller and numerous aggregates whereas W498R/C produced slightly larger aggregates but fewer in number. Live cell tracking of translational motions of these aggregates could not provide noticeable differences due to their restricted mobility. We refined our analysis through FCS by focussing into diffusional motions into infinitesimally small volumes inside the nucleoplasm where visible aggregates were absent. Interestingly enough, we obtained small diffusion coefficient and consequently longer diffusional times for R453W compared to the wt LA and W498R/C. Therefore, we interpreted this result as a contribution of motion of a somewhat bulky molecule with concomitant diffusion times. We could thus explain that the appearance of numerous visible aggregates for R453W which might have resulted from an ensemble of the highly fluctuating molecules averaged over the total nuclear space. Now the obvious question was why R453W seeded the formation of a greater number of macromolecular structures. Along this line of observation, we reasoned that the microdynamics of the Ig fold domain might hold the answer to this question. In the case of R453W, the observed RMSF from our MD simulations over 500 ns, predicted higher fluctuations resulting in higher unbalanced forces. The fluctuations occurring throughout the Ig-fold domain has been depicted by the eigenvectors. It is hypothesized that this domain exhibits faster stochastic movements leading to a higher preponderance of random self-association to form aggregates thereby lowering the probability of a stepwise steady association. Recent studies with DEAD-Box Helicase 3 X-Linked protein showed identical results with respect to protein hyperssembly based aggregation and sequestration of healthy proteins, thereby abrogating normal cell function (52). Esperante et. al. elucidated abnormal protein aggregation linked to conformational plasticity in the case of transport protein transthyretin which causes Hereditary transthyretin amyloidosis (ATTR) which is characterized by deposition of the (TTR) as amyloid fibrils in extracellular space (53). Taken together, we can opine that changes in local dynamics of a domain could indeed reflect in the molecular assembly process of a protein like lamin A. In retrospect, there is an extensive array of literature which laid out the steps for a stepwise assembly of a giant macromolecular mesh like lamina starting from lamin A dimers (54,55). Hence, we can explain thus explain the formation of broken lamina and misshapen nuclei in the case of the mutants with a more pronounced effect in the case of R453W. This can explain the severe phenotypes associated with R453W afflicted patients of autosomal dominant EDMD.

The other mutants W498R & W498C also showed fluctuations similar to R453W but lower in magnitude which was corroborated by lower RMSF values comparable to the wild type protein. Slower fluctuations would lower the rate of random association thereby producing less bulky species with higher rates of diffusion as observed from FCS. To summarize our observations, we state that the microdynamics of the Ig fold domain plays a decisive role in the stepwise lamina assembly. This is reminiscent of our observations(6) where we had predicted a partial overlap of the tail domain with a portion of the rod domain. The Ig fold thus being in the C-terminal domain could potentially exert its influence on the homooligomerization of the protein which principally occurs along the coiled-coil rod domain (residues 30-386). Therefore, any change in microdynamics of the Ig fold containing C-terminal domain could adversely affect the process of stepwise synchronized assembly into a higher order functional lamina. We conjecture for the first time that AD-EDMD and other muscular dystrophies associated with R453W and W498R/C which are largely characterized by aberrant nuclei and broken lamina could very well be caused by such abnormal dynamics of Ig fold domain.

## Acknowledgements

CM, DS, LM & M.M performed the experiments. CM, DS, LM, DB & KSG wrote the manuscript. The project was conceived by KSG. The authors declare no conflict of interest.

## Data availability statement

All representative data generated or analysed during this study are included in this submitted article and its supplementary information files. The total datasets generated during and/or analysed during the current study are available from the corresponding author on reasonable request. No mandated datasets having information or access links are used in this paper. However, the details of molecular dynamic simulation based on GROMACS and CHARMM platforms can be provided as and when required.

## Supplementary Information

**Supplementary Figure 1.**
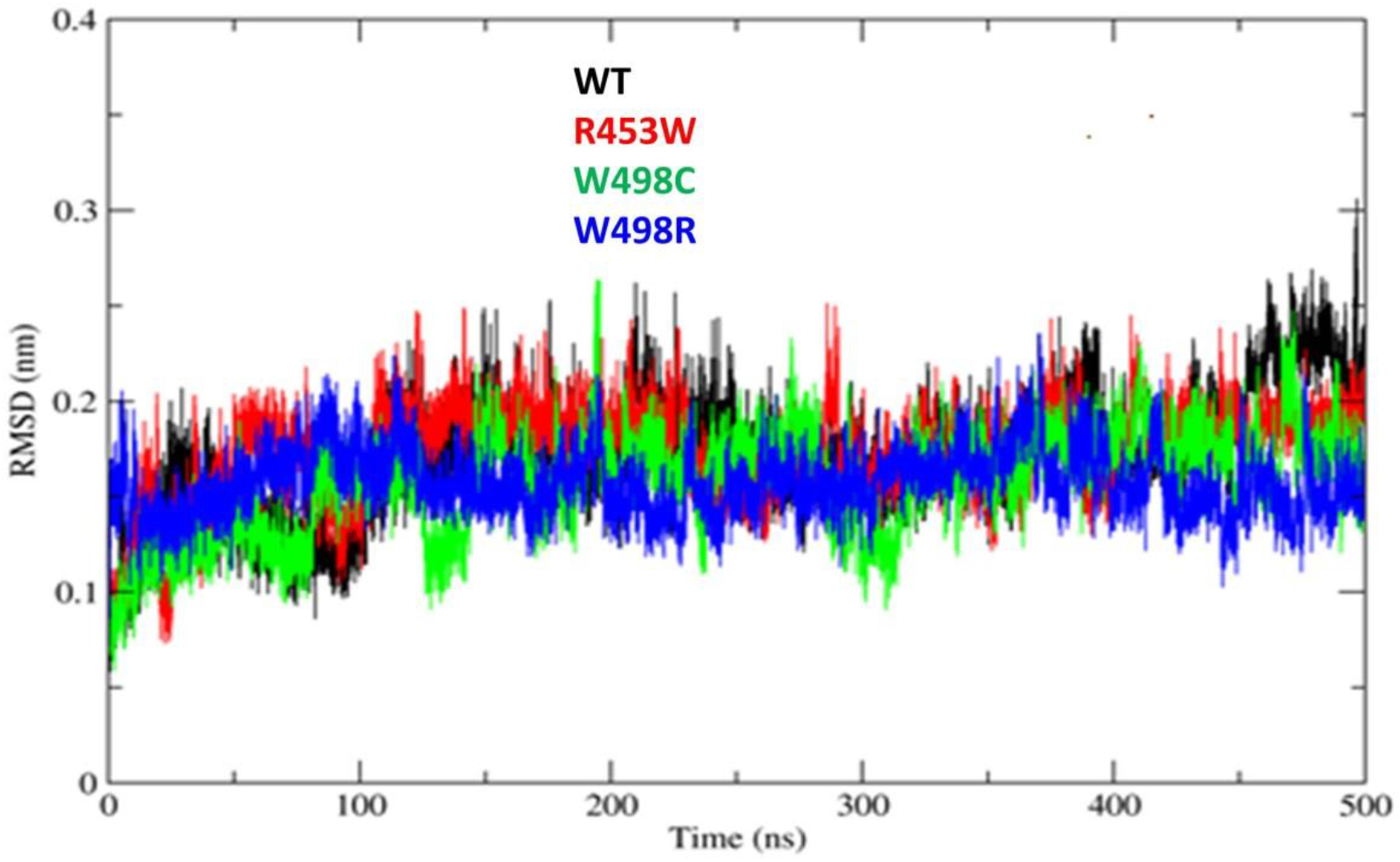
Root mean square deviation of Cα atoms of the four mutant proteins with respect to their respective energy minimized structures. C-alpha Root Mean Square Deviation (RMSD) of Ig-fold protein. All the RMSD values were calculated by using crystal structure as reference and protein Cα atoms for least fitting.

**Supplementary Figure 2.**
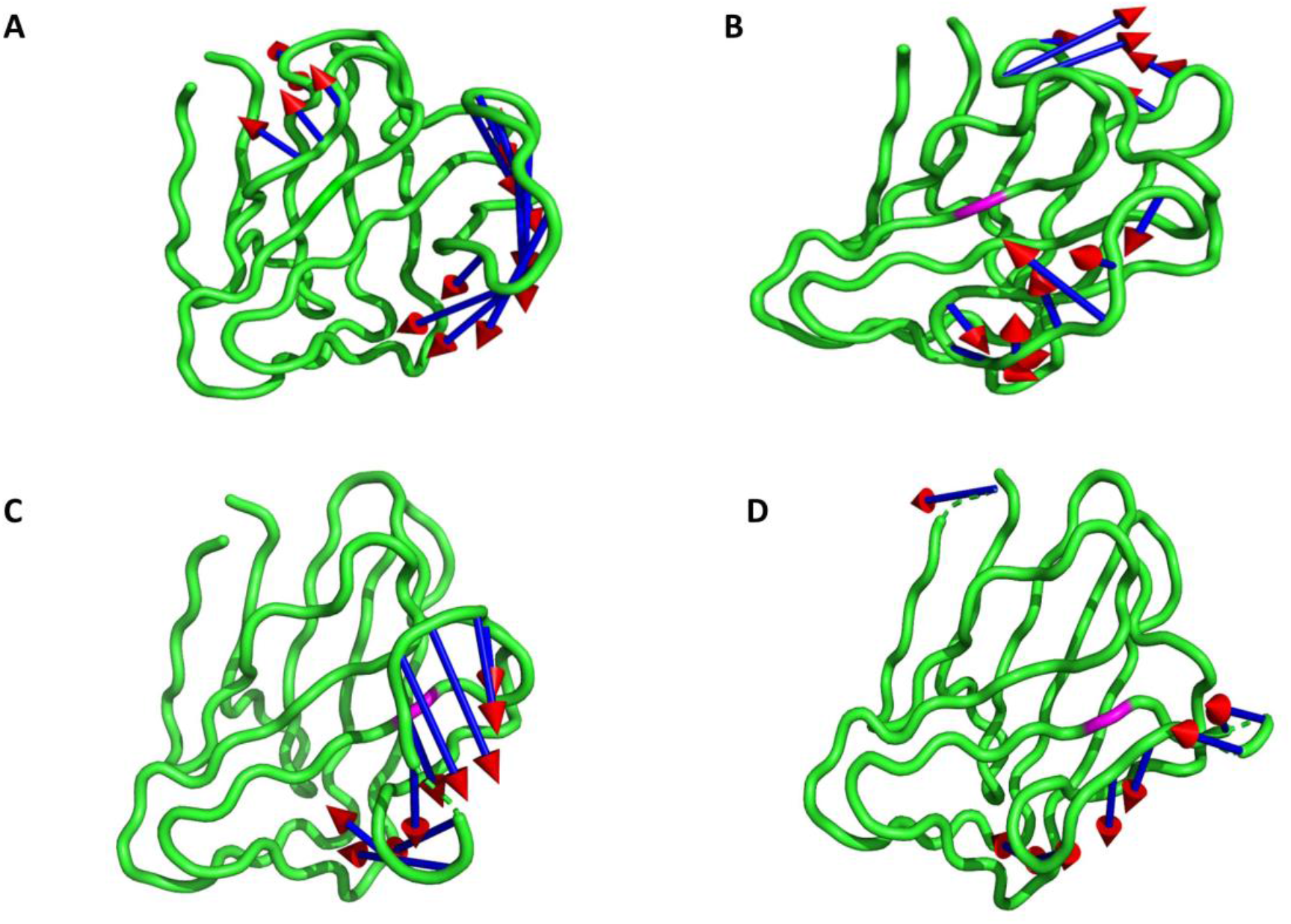
Porcupine analysis for the proteins. **(A)** WT **(B)** R453W **(C)** W498C and **(D)** W498R lamin mutants.

